# Hippocampal multi-layered RNAseq prioritizes oligodendrocyte dysfunction over immune–driven neuroinflammation in neurolupus pathogenesis

**DOI:** 10.1101/2025.10.03.679744

**Authors:** Karim Matmat, Christian Klein, Céline Keime, Ayikoé-Guy Mensah-Nyagan, Hélène Jeltsch-David

**Author notes:** Karim Matmat – Christian Klein – Céline Keime – Ayikoé-Guy Mensah-Nyagan – Hélène Jeltsch-David –.

## Abstract

Neuropsychiatric systemic lupus erythematosus (NPSLE) is a severe manifestation of lupus marked by cognitive and mood disorders, yet its hippocampal molecular underpinnings remain poorly understood. Here, we provide a region-specific transcriptomic map of the hippocampus in MRL/Lpr mice —a validated NPSLE model— compared to MRL^+/+^ controls. Bulk RNA-seq combined with integrative analyses (*e.g.* differential expression, GSEA, WGCNA, cell-type deconvolution) uncovered a robust disease-specific signature centered on oligodendrocyte dysfunction and myelination failure. Key myelin-related genes (*Mbp*, *Plp1*, *Mog*) and lineage-defining transcription factors (*Sox10*, *Nkx6-2*, *Olig2*) were repressed, while OPC markers remained unchanged, indicating a maturation blockade rather than lineage loss. Gene set enrichment highlighted widespread suppression of oligodendrocyte differentiation, axon ensheathment, and Wnt/retinoic acid signaling, alongside dysregulation of extracellular matrix components critical for axo-glial interactions. Co-expression network analysis revealed a disease-associated module enriched in myelination programs, with hub genes spanning structural, transcriptional, and adhesion-related functions. Deconvolution analysis confirmed a selective reduction of mature oligodendrocytes, contrasting with preserved neuronal populations and absence of classical astroglial or microglial activation signatures. RT-qPCR and Western blot validated the repression of myelination pathways at both mRNA and protein levels. Collectively, these findings challenge the inflammation-centric paradigm of NPSLE, revealing a cell-intrinsic vulnerability of the oligodendrocyte lineage. This conceptual shift — from immune-driven damage to impaired glial development— redefines NPSLE pathogenesis and suggests novel therapeutic avenues targeting oligodendrocyte maturation and remyelination rather than focusing solely on immunosuppression.

## Introduction

Neuropsychiatric systemic lupus erythematosus (NPSLE) represents a severe and heterogeneous complication of systemic lupus erythematosus (SLE), a chronic autoimmune disorder with a strong female predominance (female-to-male ratio ∼9:1), affecting multiple organ systems including the central nervous system (Weckerle and Niewold, 2011; Margery-Muir et al., 2017). NPSLE encompasses a wide spectrum of symptoms, *e.g.* cognitive dysfunctions, mood disturbances, psychosis, seizures, and motor deficits such as tremors or gait abnormalities (ACR ad hoc committee, 1999; Hanly et al., 2010; Jeltsch-David and Muller, 2014a). Cognitive impairment, impacting attention, memory, learning and executive function, is among the most prevalent and disabling features, occurring in up to 50% of patients (Schwartz et al., 2019; Mendelsohn et al., 2021; Ginayah et al., 2025). Despite its clinical significance, the diagnosis of NPSLE remains challenging due to its diverse clinical manifestations, the absence of specific biomarkers, and the frequent overlap with other neurological, neuropsychiatric, or metabolic conditions (Jeltsch-David and Muller, 2014a; Patel, 2024; Berndorfler et al., 2025).

Recent clinical evidence highlights structural and functional brain abnormalities in SLE, with imaging studies reporting cortical thinning, hippocampal atrophy, disrupted functional connectivity, and microstructural alterations in white matter tracts, particularly as detected by diffusion tensor imaging (Nystedt et al., 2018; Valdés Cabrera et al., 2024; Liu et al., 2025). These observations suggest that chronic immune dysregulation and inflammation in lupus may compromise brain integrity and connectivity. However, the extent and specificity of alterations, such as white matter damage, remain debated in NPSLE, and its direct link to cognitive symptoms is still not fully established (Jung et al., 2012; Wiseman et al., 2017). Moreover, human studies are inherently limited by clinical heterogeneity, treatment history, and restricted access to brain tissue, underscoring the need for controlled mechanistic investigations in experimental models (Stock et al., 2013; Patel, 2024).

In this context, the MRL/Lpr mouse stands out as the most established and extensively characterized preclinical model of NPSLE. These mice spontaneously develop systemic autoimmunity, including lymphadenopathy, high titers of autoantibodies, and progressive renal involvement marked by glomerulonephritis and proteinuria (Gulinello and Putterman, 2011; Reynolds et al., 2024). Importantly, they also exhibit a broad range of behavioral abnormalities that recapitulate key features of human neuropsychiatric syndromes, such as cognitive deficits, anxiety, depression-like behaviors, and altered locomotion (Gao et al., 2009; Muller et al., 2017; Guan and Wang, 2023; Reynolds et al., 2024; Tan et al., 2024). These phenotypes emerge in the absence of gross structural brain lesions, making this model particularly well suited to study subtle molecular and functional alterations in the central nervous system. Yet, despite the widespread use of the MRL/Lpr strain, only few studies have examined in depth the molecular underpinnings of its neuropsychiatric phenotype.

Among the brain regions affected in NPSLE, the hippocampus stands out due to its central role in learning, memory consolidation, stress regulation, and its high vulnerability to inflammatory and metabolic alterations (De Felice and Lourenco, 2015; Liu et al., 2020; Graïc et al., 2023). As one of the rare cerebral regions exhibiting persistent adult neurogenesis, the hippocampus is uniquely positioned at the interface of immune modulation, synaptic plasticity, and network-level integration (Bruel-Jungerman et al., 2007; Abrous et al., 2022; Surget and Belzung, 2022).

Notably, human imaging studies have frequently reported hippocampal volume loss and functional abnormalities in SLE patients (Appenzeller et al., 2006; Liu et al., 2025). In the MRL/Lpr model, several studies have documented hippocampal inflammation, glial activation, and reduced neurogenesis (Chesnokova et al., 2016; Bendorius et al., 2018; Nikolopoulos et al., 2023; Masanetz et al., 2025). However, to date, very few investigations have focused specifically on the transcriptomic landscape of the hippocampus in this model (Han et al., 2022), and no study has applied a region-specific, unbiased multi-layered approach to capture its global molecular alterations.

Hence, our goal was to conduct a full and systematic characterization of hippocampal transcriptional changes in the MRL/Lpr mouse model of NPSLE using an integrative, hypothesis-free RNA-seq approach. We performed bulk transcriptomic analysis of hippocampal tissue from female MRL/Lpr mice and their MRL^+/+^ controls at 17 weeks of age, a time point corresponding to the peak of both systemic disease and behavioral symptoms (Reynolds et al., 2024). Our multi-layered analytic pipeline included differential gene expression analysis, gene set enrichment (GSEA), weighted gene co-expression network analysis (WGCNA), and computational deconvolution of cell-type composition, allowing us to explore the molecular alterations of this key brain region across multiple biological dimensions.

This focused and regionally-resolved transcriptomic strategy reveals coordinated dysregulation of several pathways including myelination, glial cell development, extracellular matrix organization, and synaptic remodeling within the hippocampus. Although our study is restricted to this single structure, this focus reflects the hippocampus’ central role in cognition and mood regulation, as well as its potential as a therapeutic target whose modulation may improve neuropsychiatric symptoms. The depth and specificity of the molecular alterations uncovered emphasize the importance of considering spatial resolution when investigating the mechanisms underlying neuropsychiatric lupus. While other regions may also contribute, these findings suggest that future studies should integrate region-specific analyses to better capture the heterogeneity and complexity of brain involvement in NPSLE. By establishing a detailed molecular profile of the hippocampus in a well-validated model, our work lays the foundation for spatially-informed approaches and contributes to the identification of novel targets for therapeutic intervention.

## Methods

### Animals, experimental design, and tissue collection

Experiments were performed using 17-week-old female MRL/MpJ-Fas^lpr^/J mice (#000485 referred to as MRL/Lpr) and their congenic controls, MRL/MpJ (000486 referred to as MRL^+/+^), both obtained from The Jackson Laboratory (Bar Harbor, ME, USA). Animals were housed in ventilated cages under standard laboratory conditions (22 ± 1 °C, 12-hour light/dark cycle), with 4 mice per cage and *ad libitum* access to food and water. At 17 weeks of age, mice were anesthetized with isoflurane and euthanized by decapitation. Hippocampi were rapidly dissected on ice, snap-frozen in liquid nitrogen, and stored at −80 °C until further processing. A total of eight hippocampal samples were collected (n = 4 per group), in line with sample sizes used in similar RNA-seq studies (Han et al., 2022).

All procedures involving animals were conducted in accordance with the European Directive 2010/63/EU and the French governmental decree 2013-118. Experimental protocols were approved by the local Animal Ethics Committee (APAFIS#35144) and conducted under the supervision of authorized investigators in a certified animal facility (Faculty of Medicine, University of Strasbourg).

### RNA isolation and sequencing

Total RNA was extracted from frozen hippocampal tissue using the NucleoSpin® RNA/Protein kit (Macherey-Nagel, Germany), following the manufacturer’s instructions. RNA concentration was determined using a Qubit™ 4 Fluorometer with the Qubit™ RNA High Sensitivity Assay Kit (Thermo Fisher Scientific, USA). RNA integrity was assessed using the Agilent 2100 Bioanalyzer system (Agilent Technologies, France), with a quality cutoff set at RIN > 8 for sample inclusion. RNA-seq libraries were prepared from 800 ng of total RNA using the Illumina Stranded mRNA Prep Ligation kit and IDT for Illumina RNA UD Indexes Ligation, following the manufacturer’s protocol. After poly(A) selection and thermal fragmentation (94 °C, 8 min), libraries were amplified (12 PCR cycles), purified with SPRIselect beads, and quality-checked on a Bioanalyzer 2100. Sequencing (50 bp) was performed on a NextSeq 2000 system, targeting ∼40 million reads per sample.

### Read preprocessing, alignment, and quantification

Raw RNA-seq reads were quality-checked using *FastQC v0.11.5* to assess base quality, GC content, and adapter contamination. Adapter sequences, poly(A) tails, and low-quality bases (Phred score <20) were trimmed using *Cutadapt v4.2* (Martin, 2011) and reads shorter than 40 nucleotides were excluded. Ribosomal RNA was removed by aligning reads to rRNA sequences from RefSeq and GenBank using *Bowtie v2.3.5* (Langmead and Salzberg, 2012). Cleaned reads were then aligned to the mouse reference genome (*GRCm39*) using *STAR v2.7.10b* (Dobin et al., 2013). Gene annotation was based on the Ensembl release 111 GTF file and gene-level quantification was performed using *htseq-count v1.99.2* (Anders et al., 2015) to produce raw count matrices for downstream transcriptomic analyses.

### Transcriptomic data processing and integrative multi-layered workflow

#### Differential gene expression analysis

Differential expression analysis was conducted using *DESeq2 v1.49.4* (Love et al., 2014) using R *v4.3.2* on raw count data obtained from hippocampal RNA-seq samples, comparing MRL/Lpr and MRL^+/+^ groups. Genes with an adjusted p-value (False Discovery Rate, FDR, calculated using the Benjamini-Hochberg method) (Benjamini and Hochberg, 1995) below 0.05 and an absolute log₂ fold change greater than log₂(1.5) were considered differentially expressed (Han et al., 2022). Volcano plot was constructed using *gplot2*, *ggrepel*, and *ggExtra* to highlight both the overall distribution and a selection of biologically relevant genes. The heatmap was generated with the *pheatmap* package to visualize the expression patterns of significantly regulated genes across individual samples, with z-score normalization applied per gene and color annotations representing experimental group, sex, and gene biotype categories. To assess chromosomal distribution and large-scale transcriptional effects, a Circos plot was built using the *Circlize* package by mapping log₂ fold changes across the *mm10* genome and displaying both smoothed average profiles and individual gene-level data points, using cytoband information retrieved from the UCSC Genome Browser. Functional enrichment analysis was performed using *ShinyGO v0.77* (Ge et al., 2020) to identify overrepresented Gene Ontology categories (Biological Process, Molecular Function, Cellular Component) and Panther Pathways among the differentially expressed genes, and the top enriched terms were visualized through customized dot plots using *ggplot2*, with color encoding the statistical significance and point size reflecting gene count. Finally, protein–protein interaction networks were explored using the *STRING database v12.0*, (https://string-db.org) (Szklarczyk et al., 2023), integrating interaction evidence from experimental data, curated databases, co-expression, gene fusion, and genomic neighborhood, with a confidence score threshold set at 0.4 to retain moderately confident interactions.

#### Tissue-specific gene signature enrichment

To infer the likely tissue origin of RNA-seq samples, we performed an enrichment analysis based on tissue-specific gene signatures derived from the *Human Protein Atlas* (https://www.proteinatlas.org/) (Uhlén et al., 2015). Human gene symbols were converted to their mouse orthologs using the *biomaRt* package. Only genes uniquely associated with a single tissue and expressed in our dataset (TPM ≥ 1) were considered. For each sample, the total expression of tissue-specific genes was summed by tissue, then normalized to compute relative contributions expressed as percentages. These proportions were visualized using stacked bar plots to assess the inferred tissue composition of each sample. All steps were conducted in R *v4.3.1* using the *tidyverse* and *ggplot2* packages for data processing and visualization.

#### Gene Set Enrichment Analysis (GSEA)

GSEA was performed using the *genekitr* R package on a gene list ranked by DESeq2 Wald statistics. ENSEMBL gene identifiers were converted to ENTREZIDs using the *org.Mm.eg.db* annotation database. Gene sets were retrieved from the *MSigDB* database via *getMsigdb* function for mouse (org = “mouse”), including Gene Ontology collections: Biological Process (C5-GO-BP), Molecular Function (C5-GO-MF), and Cellular Component (C5-GO-CC). Enrichment scores were calculated using the *genGSEA* function, and gene sets with FDR-adjusted p-values < 0.05 were selected for downstream analysis. To visualize the enrichment results, dot plots were created with *ggplot2*, showing the top activated (NES > 0) and suppressed (NES < 0) gene sets based on their normalized enrichment scores. A multi-pathway enrichment visualization was also generated using *plotGSEA()* to track expression dynamics within key biological pathways such as *ensheathment of neurons*, *glial cell development*, and *mRNA processing*. A network-based representation was constructed using a custom cnetplot, linking enriched pathways to their associated genes, with node color reflecting gene fold changes. Lastly, the *upsetplot* package was used to identify genes shared across functionally related pathways.

#### Weighted Gene Co-expression Network Analysis (WGCNA)

Co-expression network analysis was conducted using the WGCNA package. Raw count data from DESeq2 were transformed using variance-stabilizing transformation (VST), and the top variable genes were selected for network construction. A soft-thresholding power (β = 10) was selected as the lowest value at which the scale-free topology fit index reached a plateau above 0.85, in line with WGCNA guidelines (Langfelder and Horvath, 2008). The resulting adjacency matrix was transformed into a topological overlap matrix (TOM), and modules of co-expressed genes were identified by hierarchical clustering and dynamic tree cut. Module eigengenes were correlated with the genotype to identify disease-associated modules. For selected modules, gene significance (GS) and module membership (kME) were computed to assess the relationship between gene–trait association and intramodular connectivity. Genes with high GS and kME (GS > 0.7 and kME > 0.8) were defined as hub genes. The relationship between kME and GS was visualized using hexbin density plots to improve readability. Gene Ontology enrichment analysis (Biological Process) was performed on each module using the *clusterProfiler* and *org.Mm.eg.db* packages following conversion of ENSEMBL gene identifiers to ENTREZIDs. For selected modules (*e.g.,* myelination), a Circos plot was constructed to visualize hub gene and their association with top 5 enriched biological terms.

#### Cell Type Deconvolution

To estimate hippocampal cell-type composition from bulk RNA-seq data, we performed a two-level deconvolution analysis using the *BisqueRNA* package in R (Jew et al., 2020). Raw count data were first normalized into TPM and subsequently log1p-transformed. Transcript lengths were retrieved from Ensembl (mmusculus_gene_ensembl) via the biomaRt package, with the median transcript length used to represent each gene for TPM calculation. Genes with low variance or mapping to mitochondrial chromosomes were excluded (Jew et al., 2020). We used the ZeiselBrainData single-cell reference (Zeisel et al., 2015), which was filtered to retain only hippocampal-derived cells, ensuring anatomical consistency. Deconvolution was performed at two hierarchical resolutions based on biologically coherent aggregation and hippocampus-specific cell-type identities (Zeisel et al., 2015). At the macro level clusters were grouped into six major cell classes including neurons (*CA1PyrInt, CA2Pyr2*, *Int1* to *Int4*, *Int5* to *Int10*, *Int13*), astrocytes (*Astro1*, *Astro2),* oligodendrocytes (*Oligo1*, *Oligo2*, *Oligo4*, *Oligo5*, *Oligo6*), microglia (*Mgl1*, *Mgl2),* vascular cells (*Vend1*, *Vend2*, *Peric*, *Vsmc*) and perivascular macrophages (*Pvm1*, *Pvm2*). At the fine level, we grouped clusters into functional subtypes based on marker expression and prior annotation (Zeisel et al., 2015), including pyramidal neurons_CA1 (*CA1PyrInt*), pyramidal neurons_CA2 (*CA2Pyr2*), pyramidal neurons_Subiculum (*SubPyr*), interneurons_Pvalb⁺ (*Int2, Int3, Int9*), interneurons_Sst⁺ (*Int1, Int2, Int4*), interneurons_Htr3a⁺ (*Int5, Int6, Int7,*), interneurons_Vip⁺ (*Int6, Int7, Int8, Int9, Int10,*), interneurons_Cck⁺ (*Int5, Int6, Int7, Int8*), interneurons_Calb2⁺ (*Int1, Int9, Int10,*), interneurons_Reln⁺ (*Int2, Int4, Int7, Int8, Int13*), interneurons_Lhx6⁺ (*Int2, Int3, Int13*), homeostatic astrocytes (*Astro1*), reactive astrocytes (*Astro2*), OPCs (*Oligo1*), immature oligodendrocytes (*Oligo2, Oligo3, Oligo4*), mature oligodendrocytes (*Oligo5, Oligo6*), homeostatic microglia (*Mgl2*), activated microglia (*Mgl1*), endothelial cells (*Vend1, Vend2*), pericytes (*Peric*), smooth muscle cells (*Vsmc*). Cell type proportions were estimated using the *ReferenceBasedDecomposition* function from BisqueRNA. Downstream analyses included stacked barplots for composition, Wilcoxon tests for group comparisons, and PCA biplots to evaluate sample separation and cell-type contributions.

### Quantitative RT-PCR Analysis

A total of 500 nanograms of RNA extracted from 4 hippocampus for each group were used for two-step reverse transcription quantitative PCR (RT-qPCR). Briefly, reverse transcription was performed using PrimeScript™ RT Master Mix, and quantitative PCR was carried out with SYBR® Premix Ex Taq™ (Takara, Kusatsu, Japan) as previously described (Matmat et al., 2023). The specific primers (Supplementary material 1) were sourced from Eurogentec (Angers, France). The cycle threshold (Ct) values were determined for each sample, and the gene expression of interest was subsequently normalized to housekeeping genes *Hprt1* and *Pol2* using the 2^ΔΔCt^ method.

### Western-blotting

Proteins were extracted using the NucleoSpin® RNA/Protein kit (Macherey-Nagel, Germany) from the same hippocampal samples used for RNA-seq and RT-qPCR, ensuring consistency across molecular assays. After protein extraction, 1% phenylmethanesulfonyl fluoride (Sigma) was added as a protease inhibitor to ensure protein stability. Protein concentrations were determined using the BCA Protein Assay Kit (Interchim) according to the manufacturer’s instructions. Equal amounts of protein (30 µg per lane) were resolved on 4–20% Mini-PROTEAN® TGX™ precast SDS-PAGE gels (Bio-Rad) and subsequently transferred to PVDF membranes (Bio-Rad). Membranes were blocked for 1 h at room temperature in 5% non-fat dry milk in TBS-T (0.1% Tween-20), incubated overnight at 4 °C with an anti-MBP antibody (Abcam #ab40390, 1:1000). Anti-β-actin antibody (Abcam #ab197277, 1:5000) used as a loading control. Appropriate HRP-conjugated secondary antibodies were applied, and detection was performed using enhanced chemiluminescence (ECL, BioRad) on a ChemiDoc imaging system (Bio-Rad). Band intensities were quantified using ImageJ *v1.53*, and MBP expression was normalized to β-actin.

### Quantification and statistical analysis

Statistical analyses were performed using GraphPad Prism (v10.5.0). Continuous variables, such as densitometry values, are presented as scatter dot plots spanning the full range of observations with a horizontal line indicating the mean ± SD. Individual data points were plotted to illustrate within-group variability and central tendency. Normality was assessed using the Shapiro–Wilk test. For normally distributed data with equal variances, group comparisons were conducted using unpaired two-tailed Student’s t-tests; when variances were unequal, Welch’s correction was applied. For non-normally distributed data, the non-parametric Mann– Whitney U test was used. A p-value < 0.05 was considered statistically significant.

## Results

### Differential gene expression analysis reveals hippocampal downregulation of myelin-associated genes in MRL/Lpr mice

To investigate hippocampal transcriptomic alterations in MRL/Lpr mice, a widely used murine model of neuropsychiatric lupus, we performed bulk RNA-seq on hippocampal samples from 17-week-old female MRL/Lpr and MRL^+/+^ mice, followed by a multi-layered analytical workflow (Fig. 1). As a preliminary quality control step, we assessed tissue identity using a reference-based deconvolution strategy grounded in transcriptomic specificity data from the Human Protein Atlas. This analysis revealed a dominant enrichment of brain-specific transcripts across all samples, with additional contributions from anatomically adjacent brain structures, such as the choroid plexus (Fig. 2A), thereby supporting the brain origin of the RNA and excluding significant contamination by peripheral tissues.

**Figure 1.**
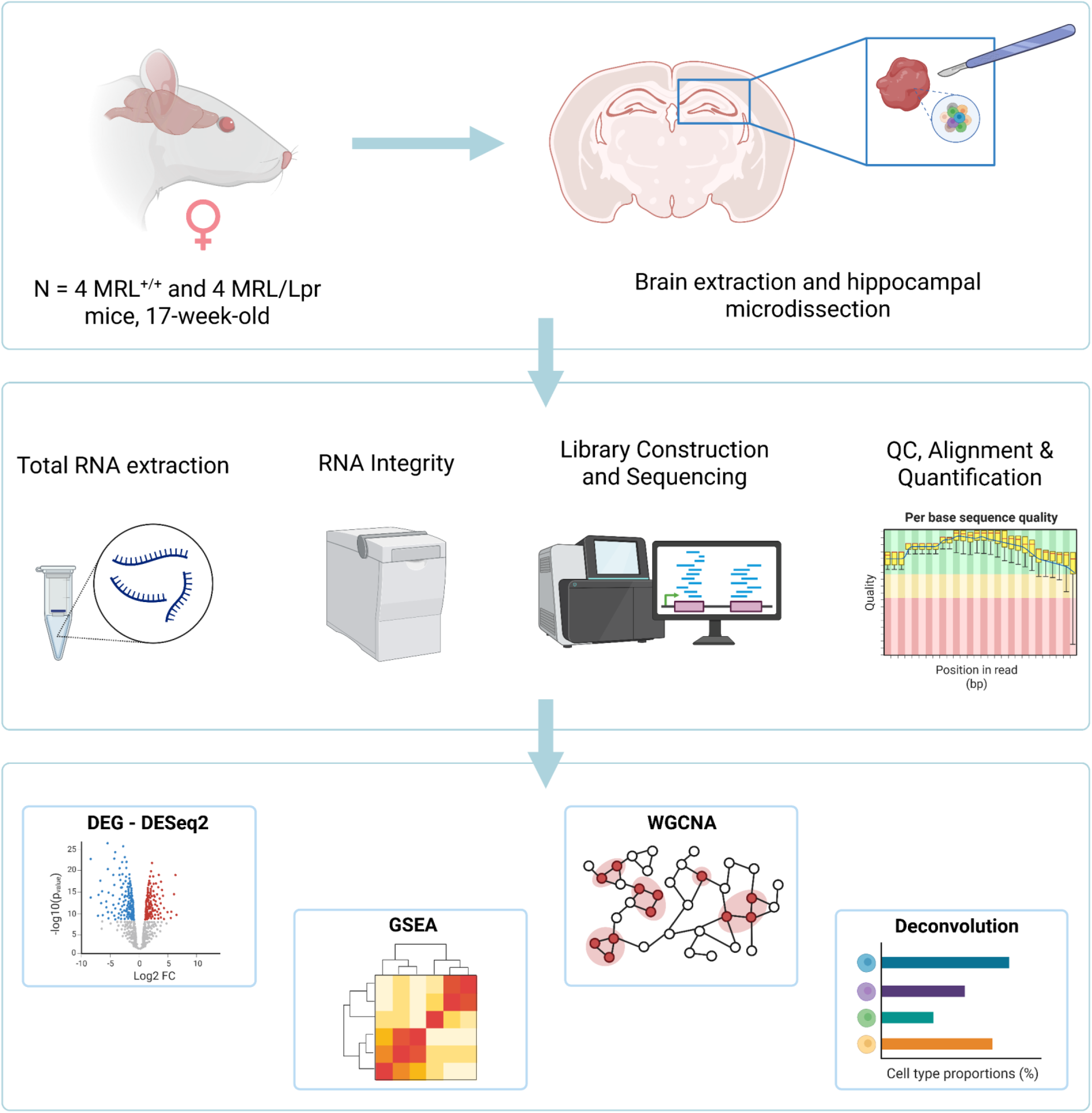
Experimental workflow and comprehensive RNA-seq analysis of the hippocampus in MRL/Lpr mice. Seventeen-week-old female MRL^+/+^ and MRL/Lpr mice (N = 4 per group) were used for hippocampal RNA-seq. The workflow included brain extraction and hippocampal microdissection, total RNA extraction, RNA integrity assessment, library construction and sequencing, followed by quality control, alignment, and gene quantification. Downstream computational analyses comprised differential gene expression analysis (DESeq2), gene set enrichment analysis (GSEA), weighted gene co-expression network analysis (WGCNA), and reference-based cell-type deconvolution (BisqueRNA). Created with BioRender.com.

**Figure 2.**
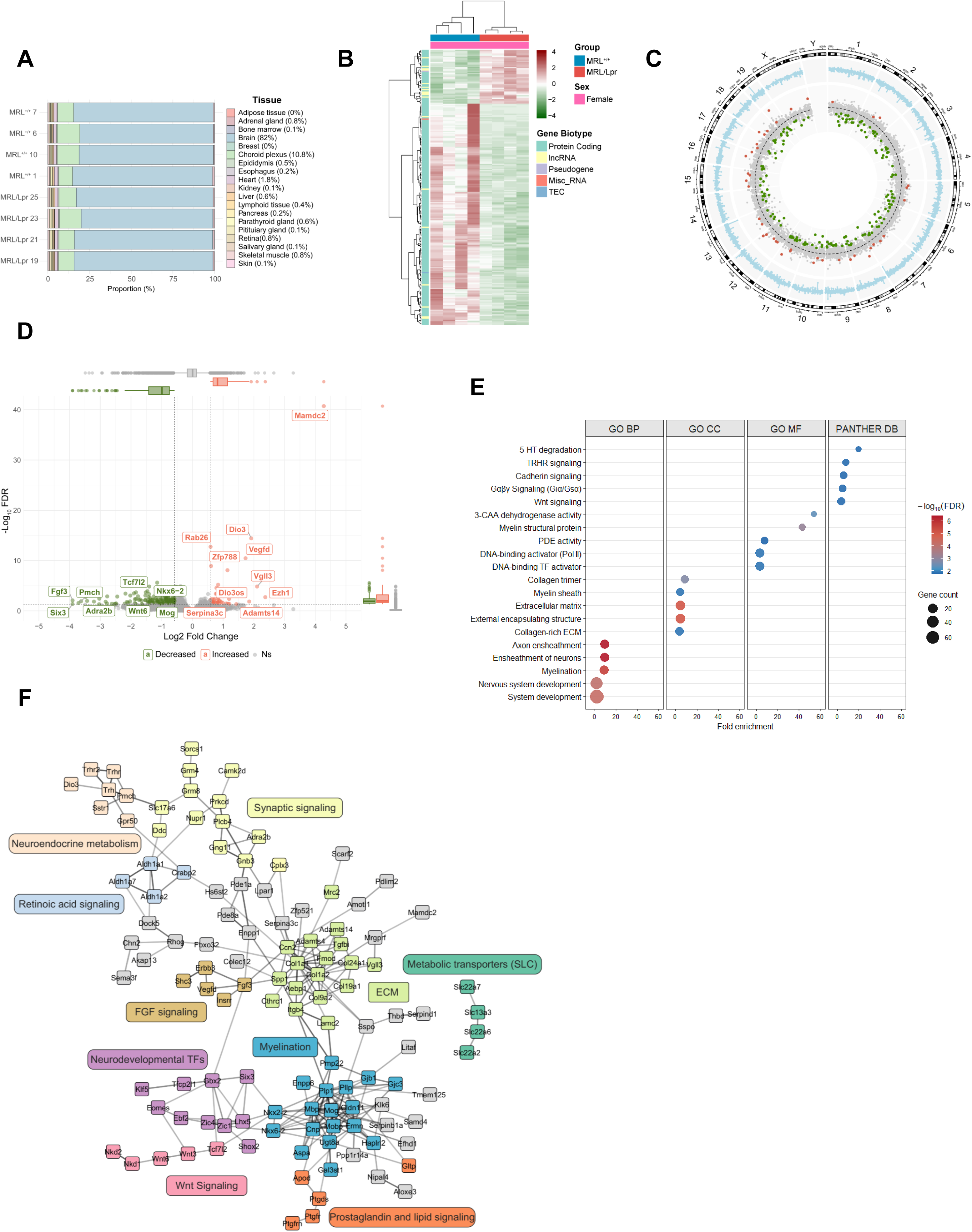
Differential gene expression and functional enrichment analyses in MRL/Lpr *vs.* MRL^+/+^ hippocampus. **(A)** Tissue origin validation based on Human Protein Atlas signatures shows a predominant brain contribution across all samples, with minor representation from choroid plexus and other tissues. **(B)** Heatmap of the 223 differentially expressed genes (FDR < 0.05; |log₂FC| > log₂(1.5)) illustrating segregation between MRL^+/+^ and MRL/Lpr groups. Rows correspond to genes and columns to samples; colors denote z-score normalized expression. Gene biotypes are annotated on the left, and sample annotations (group and sex) above. **(C)** Circos plot displaying genome-wide distribution of expression changes. Outer track (blue) shows smoothed average log₂FC across chromosomes; inner dots represent individual genes, with upregulated (red) and downregulated (green) DEGs. **(D)** Volcano plot highlighting upregulated (red) and downregulated (green) DEGs, with selected genes labeled. Boxplots above show the distribution of log₂ fold changes; boxplots on the right represent the distribution of FDR-adjusted p-values. **(E)** Top 5 functional enrichment analysis of DEGs using Gene Ontology (Biological Process, Cellular Component, Molecular Function) and PANTHER pathways; dot size indicates gene count, and color encodes –log₁₀(FDR). **(F)** Protein–protein interaction network generated from DEGs using STRING, with clusters annotated by functional categories, including myelination, Wnt signaling, extracellular matrix organization, and synaptic signaling.

Differential gene expression analysis using DESeq2 identified 223 significant differentially expressed genes (DEGs) (FDR < 0.05, fold change > 1.5), of which 180 (80.7%) were downregulated and 43 (19.3%) upregulated in MRL/Lpr mice compared to controls. The majority of DEGs (204 out of 223, 91.5%) were protein-coding genes. Hierarchical clustering of DEGs (visualized in a heatmap on Fig. 2B) separated the samples by genotype, illustrating the distinct transcriptional profiles associated with each group.

To explore the genomic landscape of expression changes, we mapped all genes across chromosomes using a Circos plot (Fig. 2C). The outermost track (blue) displays transcript expression changes in MRL/Lpr vs. MRL^+/+^. Up- and down-regulated DEGs (red and green, respectively) were distributed across all chromosomes without regional clustering, supporting a global transcriptomic shift rather than a focal genomic event. The volcano plot (Fig. 2D) highlighted the downregulation of several key genes involved in oligodendrocyte biology and myelin sheath formation, including *Mbp*, *Mog*, *Plp1*, and *Nkx6-2*, supporting the presence of myelination deficits. In contrast, upregulated genes such as *Rab26*, *Dio3*, *Mamdc2*, *Vegfd*, *Vgll3*, *Serpina3c*, *Adamts14*, and *Ezh1* are associated with diverse processes including vesicle trafficking, thyroid hormone signaling, cell adhesion, angiogenesis, inflammation, and chromatin regulation —pointing to broader changes in cellular remodeling and neuroendocrine modulation in MRL/Lpr hippocampi. Analysis of the differentially expressed genes (DEGs) through Gene Ontology (GO) and PANTHER pathway enrichment (Fig. 2E) highlighted a significant overrepresentation of pathways involved in myelination, axonal ensheathment, nervous system development, transcriptional regulation, and Wnt signaling. In addition, several neuroendocrine pathways, including those involving serotonin (5-HT) and thyrotropin-releasing hormone (TRH), were among the most enriched pathways. Processes associated with extracellular matrix organization (ECM) and remodeling also emerged as significantly enriched, suggesting transcriptional changes that may reflect structural and cellular alterations in the hippocampus of MRL/Lpr mice.

Finally, to visualize the functional relationships, we constructed a protein–protein interaction network using STRING, and displayed the graph using Cytoscape (Fig. 2F). This network revealed distinct functional clusters, with color-coded gene modules based on shared biological annotations. A prominent central blue cluster grouped genes involved in myelination, including *Mbp*, *Plp1*, and *Mog*. This module was directly connected to a pink Wnt signaling cluster and a purple neurodevelopmental transcription factor cluster, notably through the transcriptional regulators *Nkx6-2* and *Nkx2-2*, which are critical for oligodendrocyte lineage specification and differentiation. A separate green cluster, enriched for genes involved in ECM organization, was also identified. This ECM module was linked to the myelin cluster via the matrix- and adhesion-related genes *Itgb4* and *Lamc2*, suggesting potential mechanistic links between ECM remodeling and oligodendrocyte function. Given the established role of ECM components in regulating myelin sheath formation, axon–glia interactions, and oligodendrocyte maturation, this network structure reinforces the idea of coordinated disruption contributing to hippocampal dysfunction in MRL/Lpr mice.

Altogether, these findings reveal a structured transcriptional architecture centered on myelin-related genes, interconnected with signaling, developmental, and structural modules, reflecting multifaceted alterations in hippocampal cellular identity and neurodevelopmental programs in the MRL/Lpr model.

### Gene set enrichment analysis highlights coordinated suppression of myelination-related biological programs in MRL/Lpr hippocampus

To functionally contextualize the transcriptional alterations identified by DESeq2, we next performed gene set enrichment analysis (GSEA) on the ranked transcriptome. This approach allowed us to assess whether groups of functionally related genes were consistently enriched among up- or down-regulated transcripts in MRL/Lpr mice. GO Biological Process (GO:BP) enrichment analysis revealed a pronounced negative enrichment of gene sets related to oligodendrocyte differentiation, glial cell development, neuron ensheathment, and gliogenesis, consistent with widespread repression of myelination-related programs (Fig. 3A). Conversely, gene sets related to RNA metabolism, splicing, and mRNA processing were positively enriched, indicating a global shift in transcriptional regulation. GSEA enrichment curves confirmed the opposing dynamics between upregulated RNA processing pathways and downregulated glial developmental programs (Fig. 3B). Complementary enrichment analyses using GO Cellular Component (GO:CC) and Molecular Function (GO:MF) categories further supported these findings. Repressed gene sets included myelin sheath, postsynaptic membrane, and collagen-containing ECM (GO:CC, Fig. 3C), while GO:MF analysis revealed enrichment of RNA catalytic activity, RNP binding, and electron transfer among dysregulated genes (Fig. 3D). To visualize gene-level contributions to the most significantly repressed GO:BP categories, we generated a cnetplot highlighting genes associated with oligodendrocyte differentiation, glial cell development, and neuron ensheathment (Fig. 3E). These genes formed distinct yet interconnected networks, with downregulation of canonical myelin-related genes (*Mbp*, *Mog*, *Plp1*, *Aspa*, *Ugt8a)*, and several oligodendrocyte lineage transcription factors (*Nkx2-2*, *Nkx6-2*, *Sox10*, *Olig2*, *Tcf7l2*). Finally, an UpSet plot (Fig. 3F) revealed the overlap between gene sets, showing that several core genes were shared across the three myelination-related categories, further emphasizing a converging pattern of repression affecting oligodendrocyte biology.

**Figure 3.**
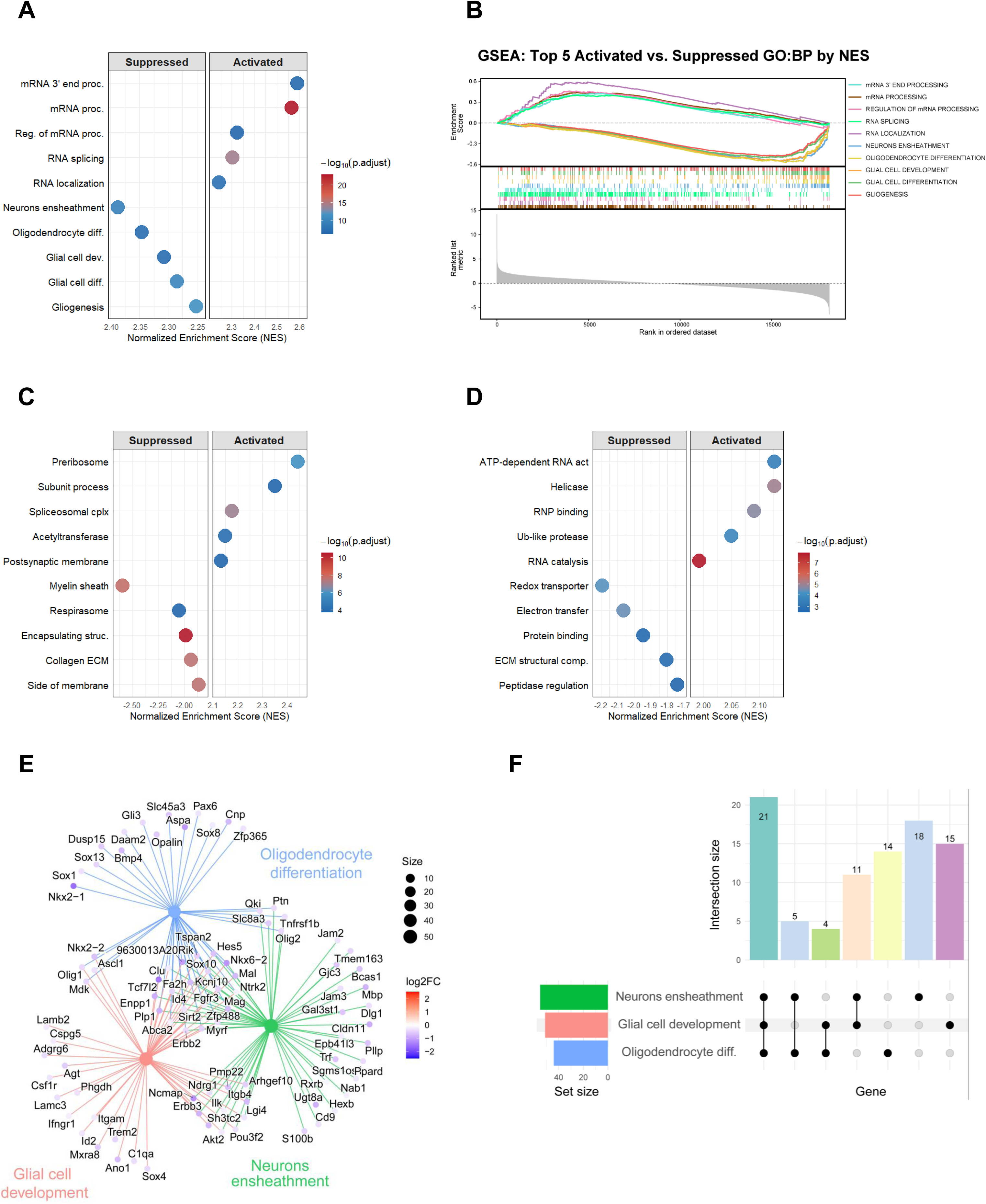
Gene Set Enrichment Analysis (GSEA) of hippocampal transcriptome in MRL/Lpr *vs.* MRL^+/+^ mice. **(A)** Top enriched GO:Biological Process (GO:BP) terms identified by GSEA, showing suppressed (left) and activated (right) pathways. Dots represent gene sets, with color indicating significance (–log₁₀ FDR), x-axis shows normalized enrichment score (NES). **(B)** GSEA enrichment curves for the top five suppressed and activated GO:BP pathways based on NES. **(C)** Top enriched GO:Cellular Component (GO:CC) terms identified by GSEA. Representation as in (A). **(D)** Top enriched GO:Molecular Function (GO:MF) terms identified by GSEA. Representation as in (A). **(E)** Network-based representation (cnetplot) linking key suppressed GO:BP categories (neurons ensheathment, glial cell development, oligodendrocyte differentiation) to their associated genes. Term nodes are sized by gene set size, while gene nodes are colored according to log₂ fold change. **(F)** UpSet plot showing intersections among genes from the three major suppressed GO:BP terms; bar height indicates intersection size, and dot connectivity indicates shared membership.

Altogether, these results highlight the presence of a coordinated transcriptional suppression of neuroglial differentiation and myelin formation programs in the hippocampus of MRL/Lpr mice, reinforcing the hypothesis of defective myelination in this model of neuropsychiatric lupus.

### Co-expression network analysis identifies a disease-associated myelination module repressed in MRL/Lpr hippocampus

To uncover coordinated transcriptional programs underlying hippocampal alterations in MRL/Lpr mice, we applied Weighted Gene Co-expression Network Analysis (WGCNA) to the full RNA-seq dataset. This approach organizes genes into modules based on shared expression patterns across samples, enabling the identification of biologically coherent gene networks.

To construct the network, we selected a soft-thresholding power of β = 10 to approximate a scale-free topology —an essential property of biological networks, in which few genes act as highly connected hubs, while most remain sparsely connected (Fig. 4A). Unsupervised clustering of transcriptome-wide expression profiles illustrated the grouping of MRL^+/+^ and MRL/Lpr samples (Fig. 4B). Gene clustering based on topological overlap —a measure of shared co-expression connectivity, resulted in the identification of multiple modules, each assigned a unique color label (Fig. 4C). Module sizes varied, with the ten largest ranging from 138 to 2,106 genes (Fig. 4D). To assess biological relevance, we correlated each module’s eigengene —a representative expression profile summarizing the module’s gene expression— with genotype. Ten modules showed significant genotype associations (Fig. 4E), and six of these (green, red, yellow, cyan, darkmagenta, and greenyellow) were prioritized for downstream analysis based on correlation strength and biological relevance.

**Figure 4.**
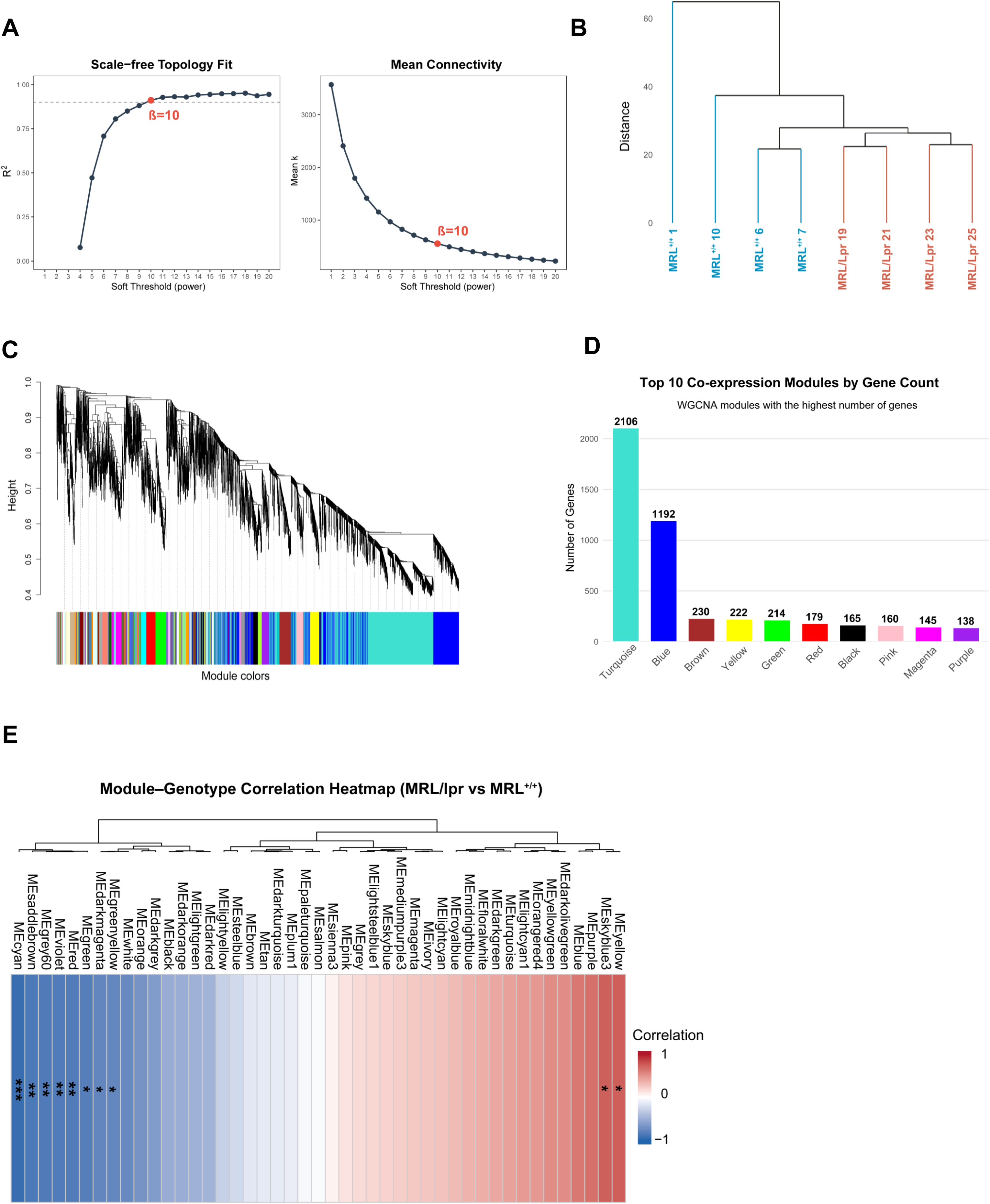
Weighted Gene Co-expression Network Analysis (WGCNA) identifies genotype-associated modules in hippocampal transcriptomes. **(A)** Scale-free topology fit index (left) and mean connectivity (right) across soft-thresholding powers. Power β = 10 was selected to approximate scale-free topology (R² ≈ 0.85), while maintaining sufficient connectivity. **(B)** Sample clustering dendrogram based on hierarchical clustering of expression profiles (blue: MRL^+/+^ and red: MRL/Lpr samples). **(C)** Gene dendrogram with assigned WGCNA modules represented by distinct colors. Modules were defined by dynamic tree cut at a minimum module size of 30 genes. **(D)** Bar chart showing the top 10 co-expression modules ranked by gene count, ranging from 2106 (turquoise) to 138 (purple). **(E)** Module–trait correlation heatmap for genotype (MRL/Lpr vs. MRL^+/+^). Each cell represents the correlation between module eigengenes and genotype, with color scale indicating correlation coefficient and asterisks marking significance (*p < 0.05; **p < 0.01; ***p < 0.001). Significance assessed from Pearson correlation using Student’s t-distribution.

We next investigated the internal architecture of these modules by comparing module membership (kME) and gene significance (GS). The kME score (also known as eigengene connectivity) quantifies how strongly a gene correlates with the module eigengene, while GS reflects the correlation between a gene’s expression and genotype. To visualize this relationship, we used hexbin density plots (Fig. 5A), which aggregate genes into hexagonal bins to improve readability in large datasets by highlighting areas of high gene density across the kME–GS plane. All six modules showed highly linear correlations between kME and GS (Pearson r ≥ 0.98), indicating that genes most central to each module were also those most strongly associated with the disease phenotype. We defined hub genes as those with both high intramodular connectivity (kME > 0.8) and strong phenotype association (GS > 0.7), representing key drivers of module function and disease relevance.

**Figure 5.**
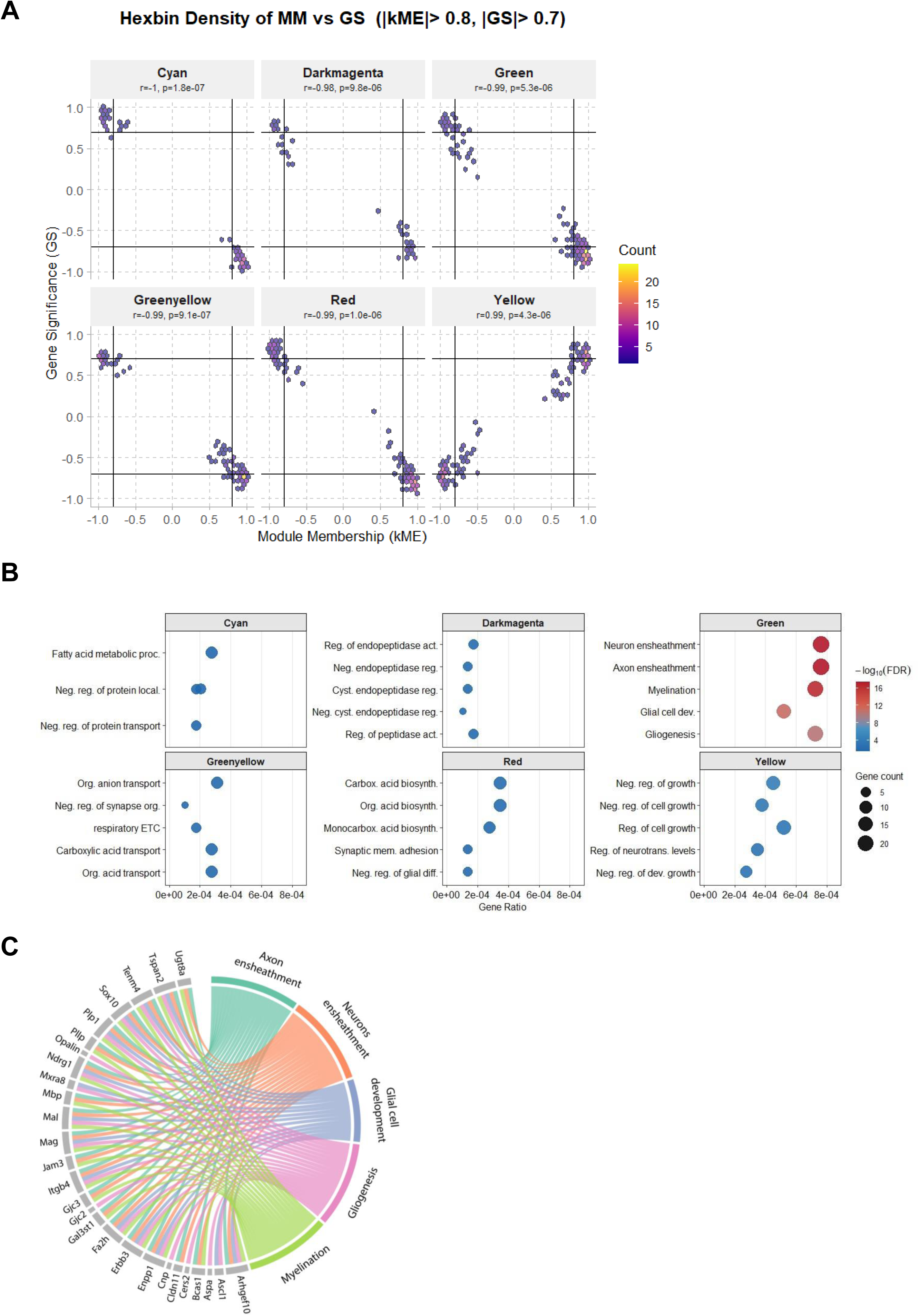
Functional characterization of hub genes in genotype-associated WGCNA modules. **(A)** Hexbin plots showing the relationship between module membership (kME) and gene significance (GS) for six genotype-associated modules (cyan, darkmagenta, green, greenyellow, red, yellow). Each hexagon represents gene density, with color scale indicating the number of genes per bin. Correlation coefficients (r) and p-values are reported for each module; vertical and horizontal lines mark hub gene thresholds (|kME| > 0.8 and |GS| > 0.7). **(B)** Functional enrichment analysis (GO:Biological Process) for each module. Dots represent enriched terms, with size indicating gene count and color representing significance (–log₁₀ FDR). **(C)** Chord diagram linking hub genes from the green module (|kME| > 0.8, |GS| > 0.7) associated with the three top enriched GO:BP terms, *e.g.* myelination, axon ensheathment, and glial cell development, identified from module enrichment analysis. Ribbons represent gene– term associations, and color denotes functional category.

Gene Ontology enrichment analyses (Fig. 5B) revealed that the green module was strongly enriched for biological processes related to myelination, axon and neuron ensheathment, glial cell development, and gliogenesis —positioning it as a central element of the transcriptional dysregulation in MRL/Lpr hippocampus. Other modules captured complementary processes, including synaptic adhesion and astrocyte differentiation (red), proteolytic regulation (darkmagenta), and mitochondrial metabolism and transport (greenyellow). To further dissect the green module, we focused on its hub genes and their contribution to the top GO:BP terms. A Circos plot (Fig. 5C) highlighted a tightly connected network of canonical myelin components (*Mbp*, *Plp1*, *Mag*, *Opalin*, *Gjc2*, *Ugt8a*), key transcription factors (*Sox10, Ascl1*), and ECM- or adhesion-related molecules (*Itgb4*, *Jam2*, *Gal3st1*, *Lama2*). These genes collectively bridge structural, developmental, and extracellular functions, suggesting they are critical effectors of glial dysfunction in MRL/Lpr mice.

Altogether, WGCNA revealed a tightly coordinated gene network centered on oligodendrocyte and myelin biology, selectively repressed in MRL/Lpr hippocampus. This module integrates transcriptional, developmental, and structural programs, offering mechanistic insight into the disrupted glial landscape that underlies hippocampal dysfunction in neuropsychiatric lupus.

### Cell-type deconvolution reveals reduced oligodendrocyte representation and specific glial alterations in MRL/Lpr hippocampus

To explore whether transcriptional differences observed in MRL/Lpr mice are accompanied by shifts in cellular composition, we performed a two-level cell-type deconvolution using a reference single-cell atlas from the mouse hippocampus (Zeisel et al., 2015). This approach included both macro-level deconvolution (major brain cell types) and fine-resolution deconvolution (cell subtype dynamics), providing a comprehensive overview of hippocampal population changes in the disease model (Fig. 6A).

**Figure 6.**
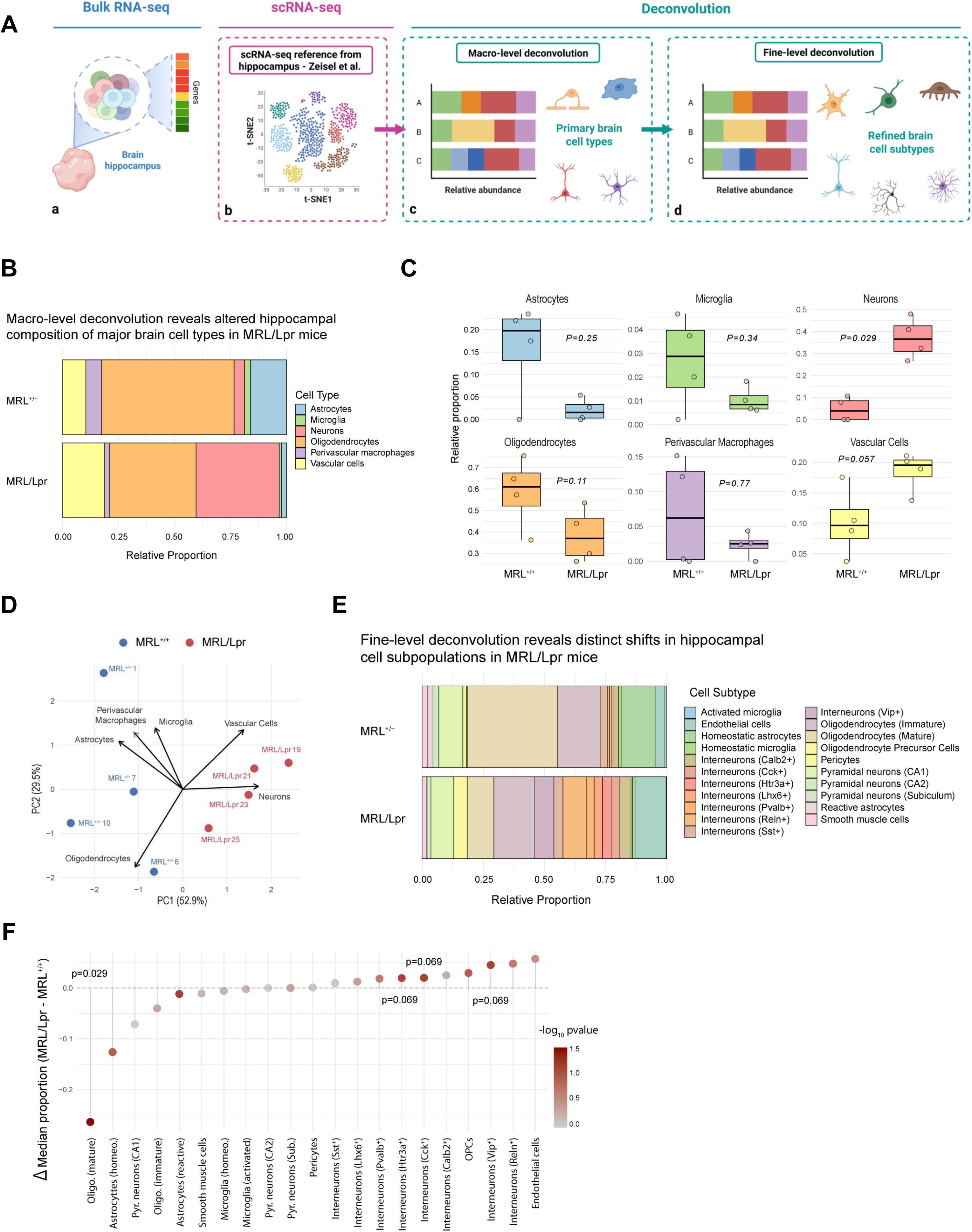
Cell-type deconvolution of hippocampal RNA-seq reveals macro- and fine-level alterations in MRL/Lpr mice. **(A)** Overview of the deconvolution workflow. Bulk RNA-seq data (a) were integrated with a single-cell reference atlas from hippocampal tissue (b) to estimate relative cell-type proportions using BisqueRNA (Zeisel et al., 2015). Two levels of analysis were performed: macro-level (primary cell classes) (c) and fine-level (specific subtypes) (d). (Created with BioRender.com). **(B)** Macro-level deconvolution showing relative proportions of major brain cell types (*e.g.* astrocytes, microglia, neurons, oligodendrocytes, perivascular macrophages, vascular cells) in MRL^+/+^ and MRL/Lpr hippocampi. **(C)** Boxplots of estimated cell-type proportions for each major population. P-values were obtained using the non-parametric Mann–Whitney U test; significant differences are indicated (p < 0.05). **(D)** Principal component analysis (PCA) based on estimated macro-level cell-type proportions. Arrows indicate loading vectors for each cell type. **(E)** Fine-level deconvolution of hippocampal subpopulations, including interneuron subtypes (*e.g.*, Vip+, Sst+), pyramidal neurons (CA1, CA2, subiculum), astrocytes (homeostatic, reactive), microglia (homeostatic, activated), and oligodendrocyte lineage cells (precursor, immature, mature). **(F)** Differential abundance analysis showing Δ median proportion (MRL/Lpr – MRL^+/+^) for each cell subtype. Dot color encodes significance level (–log₁₀ FDR), and selected p-values are displayed (Mann– Whitney U test).

At the macro level, MRL/Lpr mice exhibited signs of a neuron–glia imbalance (Fig. 6B–C). Among major cell types, only neuronal proportions were significantly increased (p = 0.029). All other populations —including oligodendrocytes (p = 0.11), astrocytes (p = 0.25), vascular cells (p = 0.057), microglia (p = 0.63), and perivascular macrophages (p = 0.74) —did not show statistically significant changes, although some exhibited non-significant trends. Principal component analysis of cell-type proportions (Fig. 6D) reinforced this interpretation. Indeed, the first principal component (PC1), accounting for 52.9% of the variance effectively separated MRL^+/+^ and MRL/Lpr samples. Importantly, neurons and oligodendrocytes contributed most strongly to this axis —highlighting their opposing dynamics and supporting the notion of a potential glia-to-neuron shift in the cellular landscape of MRL/Lpr mice.

At higher resolution, fine-level deconvolution revealed a selective reduction in mature oligodendrocyte subtypes in MRL/Lpr mice, while oligodendrocyte precursor cells (OPCs) remained largely unaffected (Fig. 6E–F). This pattern could suggest a maturation blockade rather than a global loss of the lineage, aligning with our previous findings that identified repression of key differentiation regulators such as *Nkx2-2*, *Nkx6-2*, *Ascl1* and *Sox10*.

In parallel, we observed subtype-specific changes in hippocampal interneuron composition. While global neuronal proportions were slightly elevated in MRL/Lpr mice, no significant changes were observed among pyramidal neurons from CA1, CA2, or the subiculum, which are developmentally established and not replaced in adulthood. Instead, several subsets of interneurons —notably Htr3a⁺, Cck⁺, and Vip⁺ populations— exhibited upward trends in abundance (p ≈ 0.07). These subtypes are typically associated with disinhibitory and neuromodulatory circuits. Surprisingly, and consistent with the absence of strong neuroinflammatory transcriptional signatures in our RNA-seq data, we did not observe major shifts in astrocyte or microglial activation states. Reactive vs. homeostatic astrocytes, as well as activated vs. homeostatic microglia, showed no significant differences in their proportions, suggesting that classical inflammation-driven glial remodeling may not be a dominant feature in the hippocampus of MRL/Lpr mice. It should be noted, however, that this analysis was limited to brain-resident populations and may not reflect contributions from infiltrating immune cells (*e.g.,* T cells, monocytes, B-lineage cells).

Altogether, this integrative deconvolution approach highlights a selective reduction in mature oligodendrocytes, a broader shift favoring neuronal over glial lineages, and modulation of interneuron diversity in the MRL/Lpr hippocampus. These findings corroborate our transcriptomic and network analyses, supporting a model of disrupted myelination and glial maturation as key contributors to hippocampal dysfunction in neuropsychiatric lupus.

### Molecular validation of myelination pathway repression

To validate the transcriptomic alterations observed in oligodendrocyte-related pathways, we performed targeted RT-qPCR quantification of key genes pointing successive stages of oligodendrocyte maturation: oligodendrocyte progenitor cells (OPCs, yellow), differentiation transcription factors (green), immature oligodendrocytes (purple), and mature oligodendrocytes (brown) (Fig. 7A). Quantitative analysis (Fig. 7B) revealed a significant downregulation of multiple genes involved in oligodendrocyte differentiation and function in MRL/Lpr hippocampi. While OPC markers (*Pdgfra*, *Cspg4*) remained unchanged, expression of critical transcription factors, such as *Nkx6-2*, *Olig2* and *Sox10,* was significantly decreased (*p* = 0.03, 0.01, and 0.02, respectively), suggesting impaired initiation of the differentiation program. This was accompanied by reduced levels of genes associated with (i) immature oligodendrocytes (*Cnpase*) and (ii) mature myelinating cells (*Mbp*, *Plp1*, *Mobp*), with *Mog* also trending toward downregulation (*p* = 0.07). Consistent with these transcriptional findings, western blot analysis confirmed a marked reduction in MBP protein levels in MRL/Lpr mice compared to controls (*p < 0.001*) (Fig. 7C). Together, these results corroborate the oligodendrocyte-specific modules identified in the WGCNA analysis and the suppression of myelination-related pathways observed in the GSEA.

**Figure 7.**
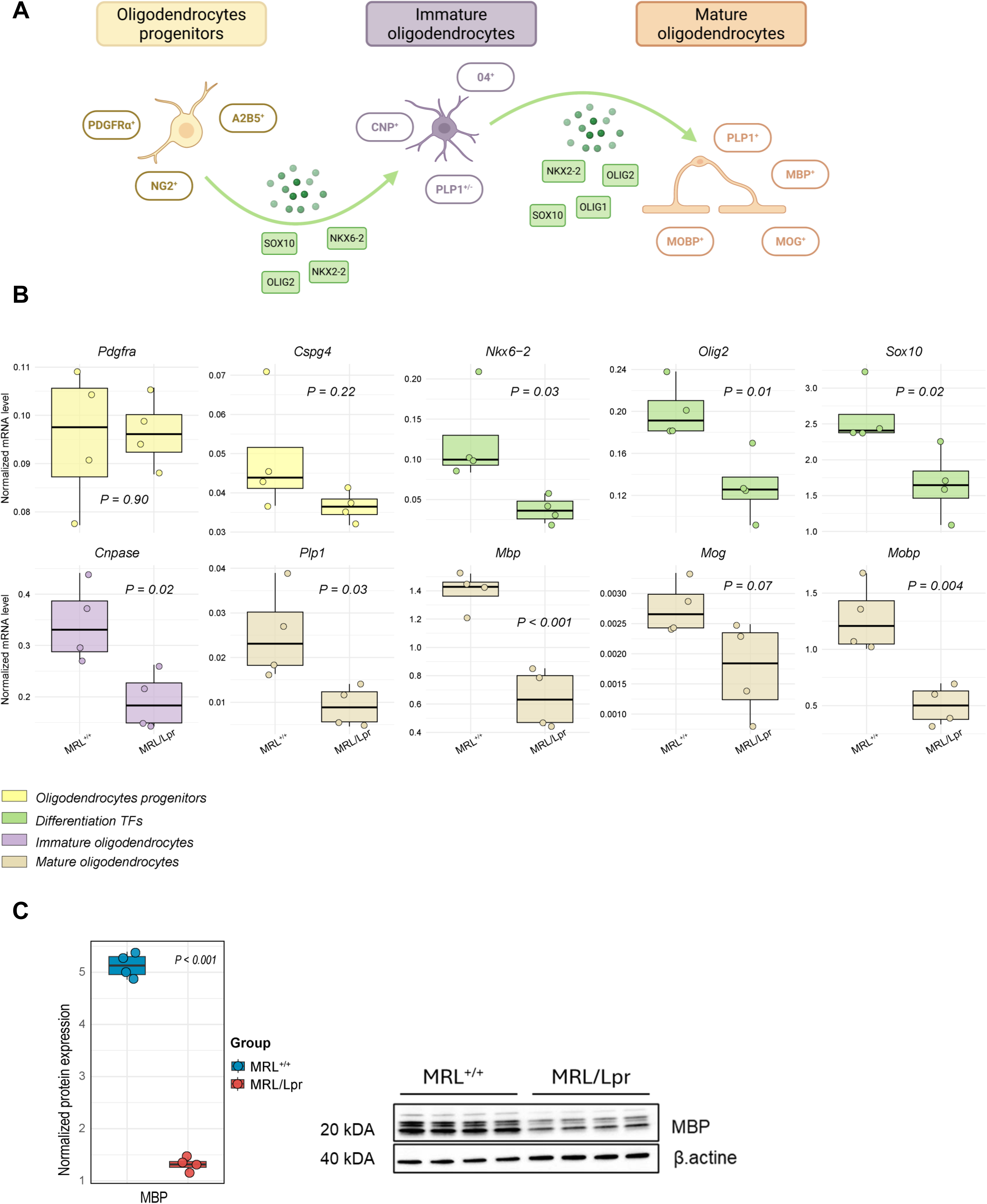
Validation of oligodendrocyte differentiation impairment in MRL/Lpr mice. **(A)** Schematic representation of the oligodendrocyte lineage, showing the transition from progenitors (PDGFRα^+^, NG2^+^, A2B5^+^) to immature (CNP^+^, PLP1^+^) and mature oligodendrocytes (PLP1^+^, MBP^+^, MOBP^+^, MOG^+^). Key transcription factors (SOX10, OLIG1, OLIG2, NKX2-2, NKX6-2) involved in differentiation are highlighted (Created with BioRender.com). **(B)** RT-qPCR analysis of genes associated with oligodendrocyte progenitors, differentiation transcription factors, immature oligodendrocytes, and mature oligodendrocytes in hippocampal samples from MRL^+/+^ and MRL/Lpr mice. Boxplots show normalized mRNA expression levels (N = 4 per group), with p-values calculated using an unpaired two-tailed Student’s t-test. **(C)** Western blot validation of MBP protein expression in hippocampal extracts. Left panel: boxplots depicting normalized protein levels (N = 4 per group), with p-values derived from an unpaired two-tailed Student’s t-test (p < 0.001). Right panel: representative immunoblot showing MBP and the loading control β-actin.

## Discussion

NPSLE continues to represent a significant clinical challenge, with the molecular mechanisms underlying its cognitive and psychiatric manifestations remaining poorly understood. Here, we applied an integrative transcriptomic strategy —encompassing differential gene expression analysis, pathway enrichment, co-expression network modeling, and cell-type deconvolution— to investigate hippocampal alterations in MRL/Lpr mice. This well-established model of spontaneous systemic autoimmunity recapitulates key features of human NPSLE, including systemic autoimmunity, behavioral changes, hippocampal structural abnormalities, and blood– brain barrier impairment (Gulinello and Putterman, 2011; Jeltsch-David and Muller, 2014a). Despite the hippocampus being highly susceptible to inflammatory and metabolic stress, its transcriptional landscape in this context remains insufficiently explored.

Our multi-layered analysis reveals a consistent repression of oligodendrocyte-related programs, with gene signatures indicating impaired differentiation across the oligodendrocyte lineage. These alterations were supported by myelination-enriched WGCNA modules and validated by RT-qPCR and western-blot. Deconvolution analyses further revealed a reduction in mature oligodendrocytes, while OPCs were relatively preserved or mildly increased, suggesting a differentiation blockade rather than a complete loss of the lineage. These results align with neuroimaging data from SLE patients showing white matter abnormalities, including microstructural alterations, and may provide a mechanistic basis for such findings (Appenzeller et al., 2008; Kozora and Filley, 2011). In contrast to other neuroinflammatory models, such as EAE or APP/PS1, which exhibit robust astroglial and microglial activation (Heneka et al., 2015; Prinz and Priller, 2017), we found no evidence of overt glial activation or pro-inflammatory gene expression in the hippocampus at this stage. Instead, our data point to subtler forms of glial dysregulation, centered on impaired oligodendrocyte maturation and myelination. These findings challenge the view that hippocampal dysfunction in NPSLE is primarily inflammation-driven and underscore the value of cell-type–resolved transcriptomic approaches for dissecting region-specific neuroimmune alterations.

Notably, one of the most prominent transcriptional alterations in our dataset involves the oligodendrocyte lineage, characterized by a coordinated downregulation of key myelin-related genes (e.g., *Mbp*, *Plp1*). While transcripts specific to mature oligodendrocytes were significantly reduced, those associated with oligodendrocyte progenitor cells (*e.g.*, *Pdgfra*, *Cspg4*) remained relatively unchanged, suggesting a disruption in lineage progression rather than a complete loss of these cells. This pattern is particularly relevant in the hippocampus, where active myelin remodeling supports cognitive functions such as learning and memory (Fields, 2015; Pan et al., 2020). Similar defects in oligodendrocyte maturation and myelin integrity have been reported in neurodegenerative and neuroinflammatory models, including APP/PS1, 5xFAD, and EAE mice (Wang et al., 2023). In MRL/Lpr mice, this repression appears selective and may reflect disrupted regulation by stage-specific transcription factors (*e.g.*, *Sox10*, *Olig2, Nkx6-2*) or extrinsic signals such as retinoic acid, Wnt/β-catenin signaling, and potentially ECM interactions. Indeed, our network analysis (Fig. 2F) and GSEA results (Fig. 3E) reveal concurrent downregulation of Wnt/β-catenin components (*Wnt6*, *Wnt3*, *Tcf7l2*) and retinoic acid pathway genes, both critical for oligodendrocyte maturation (Fancy et al., 2009; Morrison et al., 2020; Nanescu et al., 2025). While physiological Wnt activity is necessary to initiate transcriptional programs through *Tcf7l2*, which regulates transcription factors like *Sox10*, RA signaling promotes OPC differentiation and lipid synthesis essential for myelin formation. ECM components also play a vital role in oligodendrocyte migration, process extension, and myelin sheath stabilization, making ECM remodeling a key step for efficient remyelination (Lau et al., 2012). The combined suppression of these pathways likely disrupts both lineage progression and the structural cues required for effective myelination. Notably, our findings provide a mechanistic framework that complements neuroimaging studies in SLE patients (Appenzeller et al., 2006; Kozora and Filley, 2011). While such changes are often viewed as consequences of chronic CNS inflammation, their precise cellular origins remain unclear. Our data suggest that impaired oligodendrocyte maturation and myelin maintenance may represent an additional, underrecognized contributor to these structural changes. Overall, our data expand the classical view of NPSLE pathogenesis beyond inflammatory-mediated neuronal damage, emphasizing a role for glial dysregulation, particularly involving oligodendrocyte maturation in the hippocampal dysfunction associated with this disease.

One of the most striking and unexpected findings of our study is the lack of overt pro-inflammatory or glial activation signatures (*e.g.*, astrocyte, microglia) in the MRL/Lpr hippocampus. Contrary to the commonly held view of NPSLE as primarily driven by neuroinflammation, our data reveal no significant upregulation of canonical markers for astrocyte (*e.g.*, *Gfap*, *Serpina3n*, *Aqp4*) or microglial (*e.g.*, *Aif1*, *Cd68*) activation, wether at the bulk transcriptomic level or based on cell-type deconvolution estimates. This is particularly surprising given the number of studies reporting neuroinflammation in the MRL/Lpr brain, including elevated cytokine expression and glial reactivity in regions such as the cortex and hippocampus (Ballok et al., 2003; Ballok, 2007; Qiao et al., 2021; Reynolds et al., 2024). Several biological and methodological factors may explain this discrepancy. First, the hippocampus may exhibit a distinct temporal inflammatory profile, with a delayed or less intense response compared to other brain regions. Indeed, Han et al. (2022) demonstrated early upregulation of complement and inflammation-related genes in the hippocampus of MRL/Lpr mice at just 8 weeks of age —well before systemic disease onset— suggesting an early neuroinflammatory phase that precedes the oligodendrocyte and myelination deficits we observed at 17 weeks. (Han et al., 2022). Additionally, a mild systemic LPS challenge study showed that microglial activation varies considerably across brain areas, with the hippocampus exhibiting a relatively muted response (*e.g.,* fewer changes in Iba1+ and CD68+ cells) compared to regions like the habenula or cortex (Jung et al., 2022). Second, the absence of pronounced inflammatory signals does not necessarily indicate that glial cells are functionally normal. Both astrocytes and microglia can undergo substantial transcriptional or metabolic shifts without expressing classical activation markers such as *Gfap* or *Aif1*. Emerging work highlights the presence of non-canonical glial states that diverge from the traditional pro-inflammatory (A1) or neuroprotective (A2) phenotypes. For instance, astrocytes, may lose specific homeostatic functions, sch as glutamate clearance or metabolic support, while acquiring novel, context-dependent roles that may be either protective or detrimental (Escartin et al., 2021). Similarly, microglia can adopt disease-associated phenotypes not captured by canonical markers but that still profoundly influence neuronal health and synaptic remodeling (Keren-Shaul et al., 2017). It is therefore plausible that glial cells in the MRL/Lpr hippocampus enter such atypical or mixed activation states, which escape detection in bulk transcriptomic analyses and appear quiescent. Lastly, methodological constraints of bulk RNA-seq may reduce sensitivity to subtle cell-specific activation. Factors such as spatial heterogeneity, dilution effects, and compensatory mechanisms may blunt the transcriptomic footprint of inflammation. Furthermore, our cell-type deconvolution was based solely on resident cell signatures, potentially missing infiltrating immune cells. To overcome these limitations, future investigations leveraging single-cell RNA sequencing, spatial transcriptomics, or in situ hybridization will be essential to unravel these complex dynamics.

Beyond cell-type composition, our pathway-level analyses reveal a broader pattern of transcriptional dysregulation that affects more than just the oligodendrocyte lineage. Specifically, a distinct cluster related to “Synaptic signaling” includes numerous downregulated genes essential for presynaptic function, glutamatergic and monoaminergic neurotransmission, synaptic vesicle trafficking, and second messenger signaling pathways, indicating compromised synaptic integration in the hippocampus (Figs. 2F & 5B). This transcriptional repression parallels patterns seen in chronic EAE and early-stage Alzheimer’s disease, where downregulation of synaptic genes occurs before notable neuronal loss or cognitive impairment (MacKenzie-Graham et al., 2012; Berchtold et al., 2013). Notably, synaptic and plasticity-related deficits have also been documented in MRL/Lpr mice, such as impaired long-term potentiation and dendritic remodeling within the hippocampus (Šakić et al., 1998). These findings point to a possible dual-hit mechanism, where defective myelination and disrupted synaptic signaling synergize to drive network dysfunction in the MRL/Lpr hippocampus.

Interestingly, despite the widespread gene downregulation observed in MRL/Lpr hippocampi, GSEA revealed an enrichment of RNA metabolic pathways, including mRNA processing, splicing, 3′-end maturation, and RNA localization (Fig. 3A–C). This likely reflects a rebalance toward post-transcriptional control to preserve essential transcripts under overall repression. Similar phenomena have been documented in Alzheimer’s disease, where early upregulation of RNA-binding proteins correlates with disease-associated proteomic changes (Johnson et al., 2018). Moreover, aberrant splicing and altered activity of RNA-binding proteins are increasingly recognized as key factors in the pathogenesis of various neurodegenerative and autoimmune diseases (Tao et al., 2024).

To further dissect the cellular landscape of the MRL/Lpr hippocampus, we applied reference-based cell type deconvolution, which revealed shifts in the relative abundance of major cell populations. In agreement with the observed transcriptional repression of oligodendrocyte programs, we observed a marked reduction in mature oligodendrocytes, whereas OPCs remained relatively stable (Fig. 6F), supporting the idea of a differentiation blockade. Interestingly, we also detected a relative increase in neuronal proportions, especially among interneuron subtypes. While this may seem counterintuitive given the known cognitive and psychiatric deficits in MRL/Lpr mice (Gulinello and Putterman, 2011; Jeltsch-David and Muller, 2014b), it is important to note that deconvolution provides relative rather than absolute cell counts. Therefore, apparent increases in neurons likely reflect proportional shifts caused by the loss of mature oligodendrocytes, a phenomenon documented in benchmarking studies of these methods (Avila Cobos et al., 2020; Huuki-Myers et al., 2025). Future studies employing spatial transcriptomics and histological quantification will be critical to confirm whether these changes represent true alterations in cell number, size, or state. Nevertheless, we cannot exclude the possibility that the increased interneuron proportions reflect a genuine adaptive response to network instability. Enhanced recruitment or plasticity of inhibitory circuits, particularly Htr3a⁺ and Vip⁺ interneurons, has been described as a compensatory mechanism in models of synaptic dysfunction and reduced excitatory drive (Pelkey et al., 2017). Given the transcriptomic repression of synaptic signaling in the MRL/Lpr hippocampus, these interneurons dynamics may act to stabilize neural circuits during persistent glial and myelin deficits. Taken together, our findings challenge the conventional view of NPSLE as primarily an inflammatory or vascular encephalopathy. Instead, they support a model in which non-inflammatory glial dysfunction, especially impaired oligodendrocyte maturation, plays a central role in hippocampal pathology. This emerging perspective aligns with a broader conceptual shift in neuropsychiatric research, wherein glial cells —once relegated to passive support roles— are now recognized as active regulators of neuronal circuit integrity, metabolic homeostasis, and behavioral output (Allen and Lyons, 2018). In this light, the hippocampal alterations observed in MRL/Lpr mice resemble those described in other disorders where glial impairment precedes or dominates inflammatory damage, such as vanishing white matter disease (Bugiani et al., 2011), schizophrenia (Takahashi et al., 2011), or specific subtypes of major depressive disorder marked by oligodendrocyte loss (Rajkowska and Miguel-Hidalgo, 2007). These parallels suggest that cognitive and psychiatric symptoms in NPSLE may arise, at least in part, from myelination deficits and altered glial support, rather than isolated neuronal loss or immune infiltration. Although we did not detect classical astrocyte or microglial activation, our data suggest a failure of glial cells to sustain key homeostatic programs. Downregulation of Wnt signaling, lipid metabolism, and oxidative phosphorylation points to a state of glial insufficiency that compromises neuronal and myelin support independently of overt inflammation. This dysfunction may underlie the selective vulnerability of the hippocampus, a region with high synaptic remodeling, adult neurogenesis, and intense myelin turnover, making it highly dependent on astroglial and metabolic support (Fünfschilling et al., 2012). Similar processes occur in early Alzheimer’s disease, where oligodendrocyte dysfunction and myelin loss precede amyloid deposition and contribute to hippocampal atrophy (Nasrabady et al., 2018).

While our study provides a detailed molecular portrait of hippocampal dysfunction in MRL/Lpr mice, several limitations must be acknowledged. First, analysis at a single time point limits insight into the temporal progression of glial and neuronal changes. Immune activation, myelin disruption, or compensatory mechanisms may occur at earlier or later stages. Longitudinal studies combining behavioral testing and MRI will be essential to clarify whether glial repression precedes, coincides with, or follows cognitive decline. Second, bulk RNA sequencing, though comprehensive, lacks cellular and spatial resolution. Subtle alterations in discrete glial subsets, such as reactive astrocytes or disease-associated microglia, may be masked within bulk profiles. Although deconvolution partially mitigates this, single-cell or spatial transcriptomics will be crucial to resolve rare or transitional cell states and to map cell-type-specific responses with anatomical precision. Third, transcriptomic alterations identified here have not yet been directly correlated with structural or ultrastructural changes. While MRI studies report ventricular enlargement and brain volume reduction in 16-week-old MRL/Lpr mice (Bendorius et al., 2018), and Fluoro-Jade B/Golgi analyses show neuronal degeneration and dendritic loss in CA3 (Ballok et al., 2003), hippocampal myelin integrity has not been specifically examined. Thus, though supported by MRI and neuronal pathology, our findings require validation through targeted histological methods like MBP immunostaining or electron microscopy. Importantly, our data highlight the necessity of region-specific analyses. The hippocampus shows unique vulnerabilities in NPSLE and may undergo non-inflammatory, cell-intrinsic dysfunctions that whole-brain analyses fail to capture. Future research should adopt anatomically resolved approaches to avoid masking region-specific changes within heterogeneous brain tissue. Taken together, these limitations emphasize the need for longitudinal, multimodal, and spatially resolved strategies to capture the full trajectory and cellular complexity of CNS involvement in NPSLE.

From a translational standpoint, our findings suggest that targeting glial homeostasis —rather than inflammation alone— may offer therapeutic benefit in NPSLE. Pharmacological agents promoting oligodendrocyte differentiation (*e.g.*, clemastine, benztropine), enhancing mitochondrial function (*e.g.*, SIRT1 activators), or restoring key regulatory pathways like Wnt/β-catenin signaling represent promising candidates. Such approaches could be particularly relevant in patients lacking overt neuroinflammatory symptoms, where conventional immunosuppressive treatments may prove insufficient. In sum, this work lays the foundation for a revised framework to interpret hippocampal dysfunction in NPSLE focused on glial vulnerability. By shifting the emphasis from inflammation to glial dysfunction, it opens new avenues for understanding disease mechanisms and developing targeted therapies.

## Acknowledgments

We thank the GenomEast platform, member of the ‘France Génomique’ consortium (ANR-10-INBS-0009), for performing the sequencing and for the preliminary statistical analyses.

## Data Availability

RNA sequencing data have been deposited in the NCBI Gene Expression Omnibus (GEO) under accession number GSE303674. Additional datasets supporting the findings of this study, as well as all analysis scripts (including pipelines for RNA-seq processing, WGCNA, GSEA, and deconvolution), will be made available upon reasonable request to the corresponding authors.

## Author contributions

KM: Conceptualization, data curation, formal analysis, investigation, visualization, writing – original draft, writing – review & editing. CK: Investigation, writing – review & editing. CK: data curation, formal analysis, writing – review & editing. AGMN: Funding acquisition, writing – review & editing. HJD: Conceptualization, investigation, writing – original draft, writing – review & editing.

## Supplementary material

Supplementary materials : Primers_list.xlsx

Supplementary data 1 : RNASeq_analysis.xlsx

